# Midbrain microglia-integrated organoids as a next-generation tool for chronic morphine withdrawal research: A pilot study

**DOI:** 10.1101/2025.10.30.685622

**Authors:** Hyun Yi, Shue Liu, Zane R. Zeier, Keith A. Candiotti, Shuanglin Hao

**Affiliations:** Department of Anesthesiology, University of Miami Miller School of Medicine, Miami, FL33136; Department of Psychiatry & Behavioral Sciences, University of Miami Miller School of Medicine, Miami, FL33136; Miami VA Healthcare System, Miami, FL33125

**Keywords:** iPSC, microglia, midbrain organoid, morphine, Sirt3

## Abstract

In the midst of an opioid epidemic, opioid use disorder (OUD) can lead to significant clinical impairment or distress (such as opioid dependence and/or addiction) and overdose deaths. Therefore, it is significant to elucidate the exact molecular mechanisms of OUD and identify more effective therapies. Human induced pluripotent stem cell (hiPSC)-derived organoids are self-organized 3D tissues and mimic *in vivo* human organs. Increasingly, human-specific features of opioid abuse are being investigated using emerging brain organoids, which faithfully model the key functional, structural and biological complexities of neural tissues. However, a significant limitation of brain organoids is the absence of microglia which plays a fundamental role in OUD-related remodeling of neural circuits. In this pilot study, using hiPSC KOLF2.1J cell line, we developed our approach for generating both midbrain organoid and microglia and verified that our midbrain organoid was successfully integrated with microglia. We found that chronic exposure to morphine triggers key elements of immune responses including increased TNFα signaling and downregulation of sirtuin3 (Sirt3, a mitochondrial protein, preventing oxidative stress) and manganese superoxide dismutase (MnSOD, located in the mitochondrial matrix) in microglia-organoid co-cultures. Our findings suggest that immunocompetent midbrain organoids provide excellent models with which to study human-specific mechanisms of neuroinflammation and neurodegeneration. By combining disease relevance with scalability, the model system can be utilized as an effective tool for drug screening and toxicity testing.

## 1. Introduction

The ongoing opioid epidemic is one of the most challenging public health crises in the US.^1^ Opioid use disorder (OUD) induces significant impairment or distress including dependence and addiction.^2^ According to the Centers for Disease Control and Prevention, nearly five million people in the USA experience OUD, contributing to approximately 80,000 overdose deaths annually. The cellular and molecular mechanisms of drug abuse in the brain are extremely complex, targeting multiple neurotransmitter/neuropeptide systems. Oxidative stress is one of the biomarkers for neurological damage in OUD.^3-5^ Chronic morphine use can activate glia and induce neuroinflammation, releasing proinflammatory factors.^6-8^ However, the precise mechanisms underlying neuropathy in OUD have not been fully elaborated in human systems.

Human induced pluripotent stem cell (hiPSC)-derived organoids are self-organized 3D tissues, that can mimic the key functional, structural and biological complexity of an organ.^9, 10^ Differentiation of hiPSC to small brain-like structures known as brain organoids, offers an unprecedented opportunity to model human health disorders.^10-12^ Human organoids are emerging approaches to bridging the knowledge gap between animal models and human studies in many health disorders.^13, 14^ Brain organoid is an innovative technology for studying neurodevelopment and neuropathological disorders.^15-21^ Based on their human origin, iPSCs-derived brain organoids likely better match the genomic and structural features compared to animal models.^19, 21-24^ Notaras *et al*. exposed the forebrain organoids to a panel of narcotic and neuropsychiatric-risk factors and analyzed downstream alterations in transcription, proteomics, and metabolomics.^25^ Ho *et al*. reported that iPSC-derived forebrain organoids treated with opioids displayed a significant influence on transcription regulation in glial cells.^26^ Here, we successfully developed and verified the state-of-the-art midbrain microglia integrated organoids (MIO), and pretreatment with chronic morphine upregulated TNFα and down-regulated sirtuin 3 (Sirt3) and manganese superoxide dismutase (MnSOD, also named as SOD2) in the midbrain MIO.

## 2. Methods and materials

### 2.1. Human induced pluripotent stem cells (HiPSC)

The KOLF2.1J cell line was derived from the HPSI0114i-kolf2 line (from healthy normal skin tissue donated by an age 55-59, British white male).^27^ Pantazis and colleagues have deeply characterized the genomic status, functional characteristics, and differentiation potential of different iPSC lines, leading to the identification of KOLF2.1J as a lead reference cell line.^27^ Here, the parental cell line KOLF2.1J was purchased from the Jackson Laboratory (cat#JIPSC001000, The Jackson Laboratory, Bar Harbor, ME) and cultured using protocols available from the supplier.

### 2.2. Generation of midbrain organoids

Brain organoid differentiation was carried out using a standardized protocol as reported previously.^28-30^ To generate midbrain organoids in our Lab, we used STEMdiff™ Midbrain Organoid Differentiation Kits (cat# 100-1096, STEMCELL Technologies, Cambridge, MA) and STEMdiff™ Neural Organoid Maintenance Kits (cat#100-0120) following manufacture’s protocols. Briefly, KOLF2.1J cells were grown in 10 cm tissue culture plates coated with Matrigel® (cat#354277, Corning, Glendale, AZ) in 10 mL of mTeSR plus medium (cat#100-0276, STEMCELL Technologies). When iPSC were about 70 - 80% confluent and exhibited <10% of spontaneously-differentiated cells, they were dissociated into a single cell suspension. Approximately 3 x10^6^ single cells were plated into one well of an AggreWell™800 24-well plate in midbrain organoid seeding medium. The AggreWell™800 24-well plates have been treated with anti-adherence solution (cat# 07010, STEMCELL Technologies) prior to use. From day 1-5, medium was partially changed with formation medium every day to form organoids. At day 6, formatted organoids were transferred from AggreWell™800 plates to 6-well plates (ultra-low attachment) in expansion medium. Then, the medium was changed fully every 2 days until day 25. On day 25, the medium was changed with differentiation medium and changed fully every 2 days until day 43. On day 43, the medium was changed to maintenance medium and then changed fully every 2-3 days until day 80.

### 2.3. Diameter measurements of midbrain organoids

Bright field images of midbrain organoids at each stage were taken at 5x using a Leica microscope (Fluorescent M Leica/Micro CDMI 6000B). Four images of midbrain organoids were randomly selected for diameter measurements. The diameter of each organoid from each image was measured, calculated, and averaged.

### 2.4. Reverse transcription-quantitative polymerase chain reaction (RT-qPCR)

Four randomly selected midbrain organoids were collected on day 25 and 50 for gene expression analysis by RT-qPCR. Total RNA from each organoid was isolated using RNeasy mini kit (catalog #74104, Qiagen).^31^ The RNA sample was treated with DNase I on column to remove genomic DNA. One microgram of RNA was converted into cDNA using Superscript VILO master mix (catalog #11755-050, Invitrogen), and then real time PCR was performed with Fast SYBR green master mix (catalog #4385612, Applied Biosystems). Specificity of the PCR product was confirmed by running agarose gel electrophoresis. All reaction data were calculated with 2^−ΔΔCt^ values and normalized to GAPDH as an endogenous control. Target gene 2^-ΔΔCT^ (fold change) levels in the midbrain organoids was normalized to that in hiPSC cultures. To access midbrain-specific patterning, midbrain floorplate precursor markers FOXA2, LMX1A, and EN1 and more mature marginal zone dopaminergic markers PITX3, NURR1, tyrosine hydroxylase (TH), and GIRK2 primers were used for qPCR analysis. An additional neuronal marker, MAP2 primer was also used. For microglia integrated midbrain organoid culture system, TH, GFAP (astrocyte marker), and Iba1 (microglia marker), gene expression was measured by RT-qPCR (*data not shown*). For morphine withdrawal on midbrain microglia integrated organoids, TNFα, Sirt3, and SOD2 gene expression was measured by RT-qPCR. Sequence of each primer is listed as follows: GAPDH, forward 5’-GTCTCCTCTGACTTCAACAGCG-3’ and reverse 5’-ACCACCCTGTTGCTGTAGCCAA-3’; FOXA2, forward 5’-GGAACACCACTACGCCTTCAAC-3’ and reverse 5’-AGTGCATCACCTGTTCGTAGGC-3’; EN1, forward 5’-GTGGTCAAAACTGACTCGCAGC-3’ and reverse 5’-CCGCTTGTCCTCCTTCTCGTTC-3’; LMX1A, forward 5’-CATCGAGCAGAGTGTCTACAGC-3’ and reverse 5’-TGTCGTCGCTATCCAGGTCATG-3’; PITX3, forward 5’-AGGAGATCGCCGTGTGGACCA-3’ and reverse 5’-CCGCGAAGCTGCCTTTGCATAG-3’; NURR1, forward 5’-AAACTGCCCAGTGGACAAGCGT-3’ and reverse 5’-GCTCTTCGGTTTCGAGGGCAAA-3’; TH, forward 5’-GCTGGACAAGTGTCATCACCTG-3’ and reverse 5’-CCTGTACTGGAAGGCGATCTCA-3’; GIRK2, forward 5’-CCAACAGAGTCCTTTCTGGGAG-3’ and reverse 5’-CCACAGGATCTCACTGGTGATG-3’; MAP2, forward 5’-AGGCTGTAGCAGTCCTGAAAGG-3’ and reverse 5’-CTTCCTCCACTGTGACAGTCTG-3’; GFAP, forward 5’-CTGGAGAGGAAGATTGAGTCGC-3’ and reverse 5’-ACGTCAAGCTCCACATGGACCT-3’; Iba1, forward 5’-GCTATGAGCCAAACCAGGGA-3’ and reverse 5’-GGATCGTCTAGGAATTGCTTGTT-3’; TNFα, forward 5’-GCTGCACTTTGGAGTGATCG-3’ and reverse 5’-GCTTGAGGGTTTGCTACAACA-3’; Sirt3, forward 5’-GCGGCAGGGACGATTATTA -3’ and reverse 5’-TGCCTCCACTTCCAACAACA-3’; SOD2, forward 5’-CGTTGGCCAAGGGAGATGTT-3’ and reverse 5’-CACGTTTGATGGCTTCCAGC-3’.

### 2.5. Generation of Hematopoietic Progenitor Cells (HPCs)

To generate HPCs, KOLF2.1J iPSCs were differentiated using the STEMdiff™ Hematopoietic Kit (cat#05310, STEMCELL Technologies) following manufacturer’s protocol. One day prior to day 0, iPSCs were seeded as small 100-200 μm aggregates in Matrigel®-coated 12-well plates with mTeSR plus medium. When the number of iPSC colonies was within 4-10/cm^2^, on day 0, mTeSR plus medium was replaced with Medium A to induce iPSC toward the mesoderm-like lineage. On day 2, half fresh medium A was changed. On day 3, the medium was changed to Medium B to promote further differentiation into hematopoietic cells. Then, on day 5, 7, and 10, half fresh medium B was changed. On day 12, HPCs floating in the culture supernatant, were collected for microglia differentiation for the assessment of differentiation efficiency by flow cytometry.

### 2.6. Assessment of HPC differentiation efficiency

To assess differentiation efficiency of HPCs, flow cytometry was performed on HPCs following 12 days of hematopoietic differentiation. For the expression of HPCs surface markers, anti-human CD43-APC, anti-human CD45-FITC, and anti-human CD34-PE antibodies were used. To distinguish between live and dead cells, Live/Dead Fixable Aqua Dead Cell stain kit (cat# L34965, ThermoFisher, Waltham, MA) was used. On day 12, HPCs were collected and resuspended in 1ml PBS to a concentration of 0.5x10^6 cells/ml. Six samples were prepared as follows: 1. CD43-APC alone, 2.CD45-FITC alone, 3. CD34-PE alone, 4. CD43-APC+CD45-FITC+CD34-PE + Aqua dye, 5. Aqua dye alone, and 6. Unstained cells. One µl of the reconstituted fluorescent aqua dye was added to 1 ml of cell suspension. Cells were incubated on ice for 30 minutes, protected from light. Cells were washed once with 1ml FACS buffer (PBS with 2% FBS) and then resuspend with 100 µl FACS buffer.

For each corresponding sample, 5 µl of antibodies were added and incubated at 4°C for 30 minutes in the dark. Cells were washed with PBS and then fixed in 100 µl of 37% formaldehyde at room temperature for 15 minutes. Cells were washed again with 1ml FACS buffer and resuspended with 500µl FACS buffer. Cells were filtered through 5ml falcon round-bottom tubes with a strainer and processes in the Flow Cytometry Shared Resource (FCSR) at the University of Miami. Individual samples were passed through the BD Biosciences FACS Canto-II flow cytometer, equipped with 3 lasers (405nm violet laser, 488nm blue laser and 640nm red laser), and flow data were generated and analyzed with BD FACSDIVA 9.0 software. Doublets and background noise were eliminated, and only live cells were counted for data analysis. Antibody information is listed as follows: mouse anti-human CD43, clone CD43-10G7, APC (1:50, 60085AZ.1, STEMCELL Technologies, Cambridge, MA); mouse anti-human CD45, clone H130, FITC (1:50, 60018FI.1, STEMCELL Technologies, Cambridge, MA); mouse anti-human CD34, clone 581, PE (1:50, 60013PE.1, STEMCELL Technologies, Cambridge, MA).

### 2.7. Generation of hiPSC-derived microglia

To generate microglia from iPSC-derived HPCs, the STEMdiff™ Microglia Differentiation Kits (cat#100-0019, STEMCELL Tech, Cambridge, MA) and STEMdiff™ Microglia Maturation Kits (cat#100-0020, STEMCELL Tech, Cambridge, MA) was used following the manufacturer’s protocol. After HPCs were > 90% CD43^+^ at 12 days, microglia differentiation was performed as follows: on day 0, 2 x 10^5^ HPCs were seeded into one well of a Matrigel®-coated 6 well plate containing 2ml of microglia differentiation medium. Cells were cultured at 37°C with 5% CO_2_ incubator. Half-medium was added every other day until day 12. On day 12, suspended and semi-suspended cells were collected and re-plated on to one well of Matrigel®-coated 6-well plates. Half-medium was added every other day until day 24. On day 24, entire suspended and semi-suspended cells were collected and co-cultured with midbrain organoids.

### 2.8. Assessment of microglia differentiation efficiency

After 24 days of microglia differentiation from HPCs, 2 x 10^4^ cells were plated on Poly-L-lysine coated 8-well chamber slides (cat#80824, Fitchburg, WI) in 200µl of microglia maturation medium (cat#100-0020, STEMCELL Tech, Cambridge, MA) for 2 days, and then cells were fixed with 4% PFA for 30 minutes at room temperature. Slides were incubated in 200µl blocking buffer (10% goat serum + 0.3% Triton-X in PBS) for 30 minutes at room temperature. Slides were then incubated with primary antibodies, Iba1, CD11b(OX-42), and CD45-FITC in 200 µL of 2% goat serum with 0.3% Triton-X in PBS for 1 hour at room temperature. Slides were washed twice with PBS and then incubated with secondary antibodies in 200µl of 2% goat serum with 0.3% Triton-X in PBS and for 30 minutes at room temperature. Slides were subsequently washed twice with PBS and counterstained with 200µl of 100ng/mL DAPI (cat#D9542, Sigma, St. Louis, MO) for 10 minutes at room temperature then mounted with Fluoromount-G™ slide mounting medium (cat#17984-25, EMS, Hatfield, PA). Fluorescence images were captured using a fluorescent microscope with a 20x objective. Four images were randomly selected for counting Iba1^+^ cells and CD11b^+^CD45^+^ cells with DAPI, and positive cells were calculated as a percentage of the total number of cells. Cultures of microglia, as defined by a high percentage (>90%) of CD11b^+^CD45^+^ and Iba1^+^ cells were used to produce microglia-midbrain organoid co-cultures. Primary antibody information is listed as follows: rabbit anti-human Iba1 (1:2000, 019-19741, Fujifilm Wako Pure Chemical corporation, Richmond, VA); mouse anti-human CD11b (1:100, CBL1512, Millipore, Burlington, MA); mouse anti-human CD45-FITC (1:50, 60018, STEMCELL Technologies, Cambridge, MA). Secondary antibody information is listed as follows: donkey anti-rabbit Alexa Fluor 594 (1:1000, A21207, ThermoFisher Scientific, Waltham, MA); donkey anti-mouse Alexa Fluor 594 (1:1000, A21203, ThermoFisher Scientific).

### 2.9. Immunohistochemistry of midbrain organoid

Midbrain organoids were fixed with 4% paraformaldehyde (PFA) overnight at 4 °C. Then, the organoids were cryoprotected with a 30% sucrose solution in PBS overnight at 4 °C, and placed in plastic molds with O.C.T. compound for 30 minutes at RT before freezing with liquid nitrogen. Frozen tissue blocks were sectioned using a cryostat and slices transferred to glass microscope slides. After drying, the tissue sections were hybridized with primary antibodies overnight at 4 °C, then washed and incubated with secondary antibodies for 30 minutes at room temperature, and counterstained with DAPI for 15 minutes at room temperature. Fluorescence images were captured using a Leica microscope. Antibodies used for accessing midbrain organoid were forkhead box protein A2 (FoxA2), floorplate precursor marker at day 25, microtubule-associated protein 2 (MAP2), G protein-coupled inwardly rectifying potassium 2 (GIRK2, also known as KCNJ6), and tyrosine hydroxylase (TH) at day 50 for neuronal marker and more mature marginal zone dopaminergic marker. More complete immunohistochemistry with Iba 1, GFAP, NeuN (neuronal marker), MAP2 and TH, and MnSOD expression was conducted to access characterization of midbrain microglia integrated organoids at day 88. Primary antibody information is listed as follows: rabbit anti-human FoxA2 (1:200, D56D6XP, Cell Signaling, Danver, MA); mouse anti-human MAP2 (1:500, MA5-47450, Invitrogen, Waltham, MA); mouse anti-human GIRK2 (1:500, 21647-1-AP, ThermoFisher Scientific, Waltham, MA); rabbit anti-human TH (1:200, MA5-45023, Invitrogen, Waltham, MA); rabbit anti-human Iba1 (1:2000, 019-19741, Fujifilm Wako Pure Chemical corporation, Richmond, VA); mouse anti-human GFAP (1:2000, G3893, Sigma, St. Louis, MO); rabbit anti-human NeuN (1:2000, ABN78, Millipore, Burlington, MA); rabbit anti-human MnSOD (1:200, ab137037, Abcam, Cambridge, MA). Secondary antibody information is listed as follows: goat anti-rabbit Alexa Fluor 594 (1:2000, A11037, ThermoFisher Scientific, Waltham, MA); goat anti-mouse Alexa Fluor 594 (1:2000, A11005, ThermoFisher Scientific.); goat anti-rabbit Alexa Fluor 488 (1:2000, A11008, ThermoFisher Scientific.); donkey anti-mouse Alexa Fluor 488 (1:2000, A21202, ThermoFisher Scientific).

### 2.10. Assessment of microglia integration within midbrain organoids

Midbrain organoids (80 days of differentiation) were co-cultured with differentiated microglia at a ratio of 2.5x10^5^ microglia/organoid in a volume of 0.5ml co-culture medium (0.5ml organoid maintenance medium with 2 ul of microglia supplement B (cat#100-0023, STEMCELL Tech), following the manufacturer’s instructions. Co-cultures were incubated for 7 days to allow for microglia integration into midbrain organoids. Subsequently, immunohistochemistry and RT-qPCR were performed to confirm the presence of microglia within midbrain organoids. Antibodies used for IHC are Iba1, NeuN, TH, and GFAP. Primers used for RT-qPCR are TH, GFAP, and Iba1 (*data not shown*).

### 2.11. Spontaneous morphine withdrawal model

After 7 days, microglia-integrated organoids (MIO) were treated with morphine (Supported by the National Institute on Drug Abuse) or saline as a control for 5 days at increasing concentrations of morphine (10μM, 20μM, 30μM, 40μM and 50μM) each day. After 5 days of morphine treatment, fresh co-culture medium was replaced without morphine for 7 days. Morphine treated MIO were then collected for downstream neurochemical analyses.

### 2.12. Statistical analysis

Statistical analyses were performed using GraphPad Prism 10.4 (GraphPad Software, San Diego, CA). Two-tailed *t*-test or one-way ANOVA with Holm-Sidak’s multiple comparisons test was used for neurochemical studies. The data are presented as a mean with standard error of the mean (mean ± SEM), with *p*-value of <0.05 considered statistically significant.

## 3. Results

### 3.1. Authentication of human midbrain organoids

Midbrain organoids were generated using KOLF2.1J iPSC and the STEMdiff™ midbrain Organoid Kit (see methods and materials). The KOLF2.1J line is derived from the HPSI0114i-kolf2 line (reprogrammed from *healthy normal skin tissue* donated by an age 55-59, British white male)^27^ and has been used extensively to study neurological disorders.^32, 33^ Pantazis and colleagues have deeply characterized the genomic status, functional characteristics, and differentiation potential of different iPSC lines, leading to the identification of KOLF2.1J as a lead reference cell line.^27^ Dobner and colleagues assessed mtDNA integrity, and reported that KOLF2.1J mtDNA integrity was intact and is still preserved in the commercially distributed cell line, and that the basal KOLF2.1J metabolome profile was similar to that of the two commercially available iPSC lines IMR90 and iPSC12, further validating KOLF2.1J as a reference iPSC line.^34^ Ryan reported that large structural variants in KOLF2.1J are unlikely to compromise neurological disease.^35^

The expansion and differentiation of midbrain organoids was assessed by brightfield microscopy at 5x magnification and the diameter computed at various time points for 80 days. Images were randomly selected for diameter measurement from each time point (**Figure 1A**). The size of our midbrain organoid was consistent with expected results reported by Stemcell Technologies Inc (**Figure 1B**).

**Figure 1.**
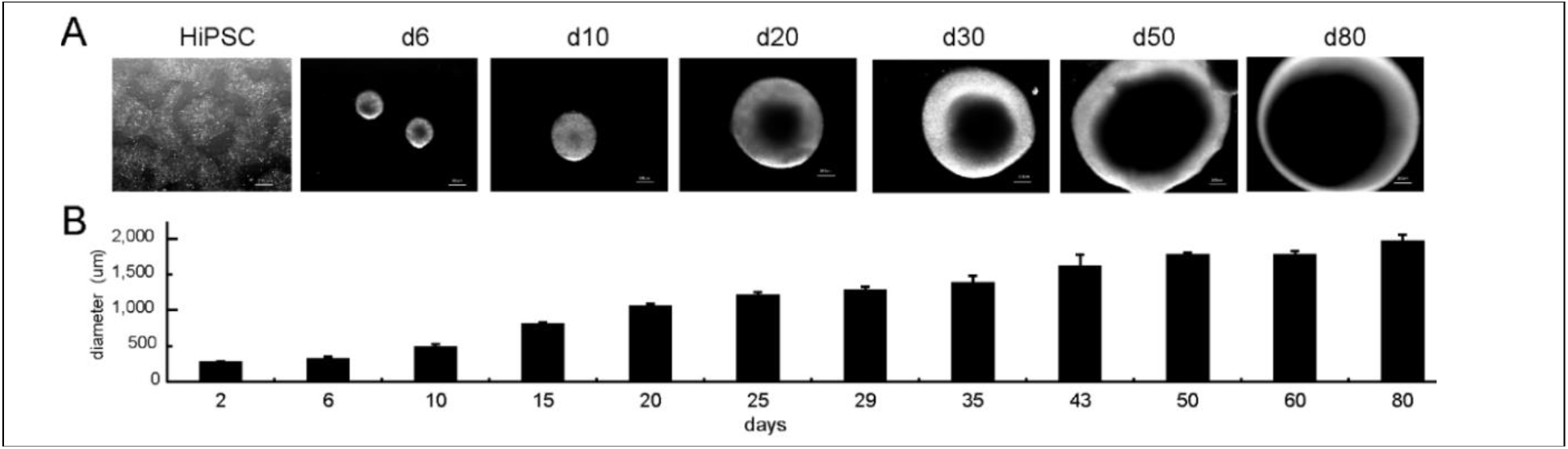
Expansion and differentiation of midbrain organoids from hiPSC. (**A**) Representative images of hiPSC culture and midbrain organoids at different time points, scale bar = 200μm. (**B**) Quantification of midbrain organoid diameter from day 2-80 of differentiation, n=4.

### 3.2. The expression of key midbrain markers within organoids was confirmed using immunofluorescence

The efficiency of midbrain organoid generation was assessed by immunohistochemistry. Immunofluorescence corresponding to the expression of the key midbrain floorplate precursor marker forkhead box protein A2 (FoxA2) in day-25 organoids is shown (**Figure 2A**), and the expression of FoxA2, a pan-neuronal marker, microtubule-associated protein 2 (MAP2), and dopaminergic markers, G protein-coupled inwardly rectifying potassium 2 (GIRK2, also known as KCNJ6), and tyrosine hydroxylase (TH) in day-50 organoids (**Figure 2B**).

**Figure 2A.**
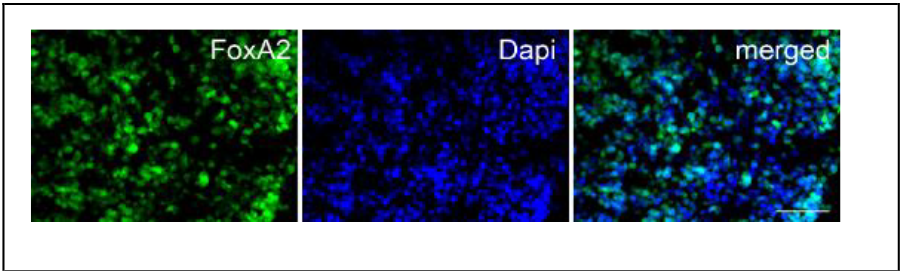
The immunofluorescence staining shows the expression of key midbrain floorplate precursor marker FoxA2 at day 25 organoids, scale bar, 50um.

**Figure 2B.**
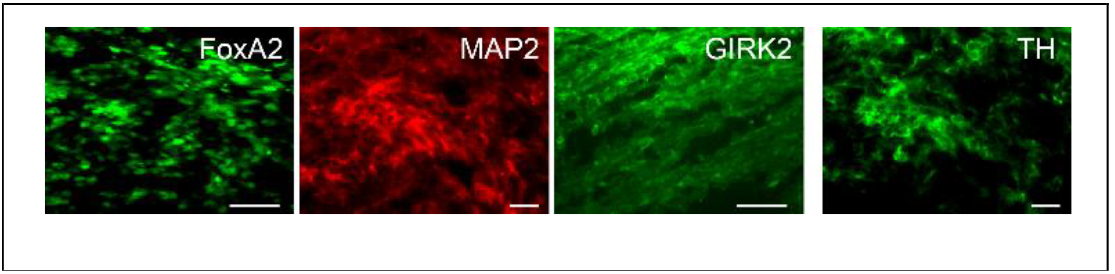
The immunofluorescence staining shows the expression of midbrain precursor marker FOXA2, neuronal marker MAP2, and dopaminergic markers GIRK2 and TH at day-50 organoids, scale bar, 50um

### 3.3. The expression of key midbrain genes in the organoids was detected using RT-qPCR

Four randomly selected midbrain organoids were collected on day 25 and 50, and analyzed by RT-qPCR as described previously.^36^ To access midbrain key markers of brain-specific patterning, we examined mRNA expression of midbrain floorplate precursor markers FOXA2, EN1, and LMX1A on day 25, and more mature markers including pituitary homeobox 3 (PITX3), nuclear receptor-related 1(NURR1), TH, GIRK2, and MAP2 on day 50 using RT-qPCR. Expression of these markers is consistent with the successful expansion and maturation of midbrain organoids that display characteristic features of midbrain neuronal fate specification and differentiation (**Figure 3**).

**Figure 3.**
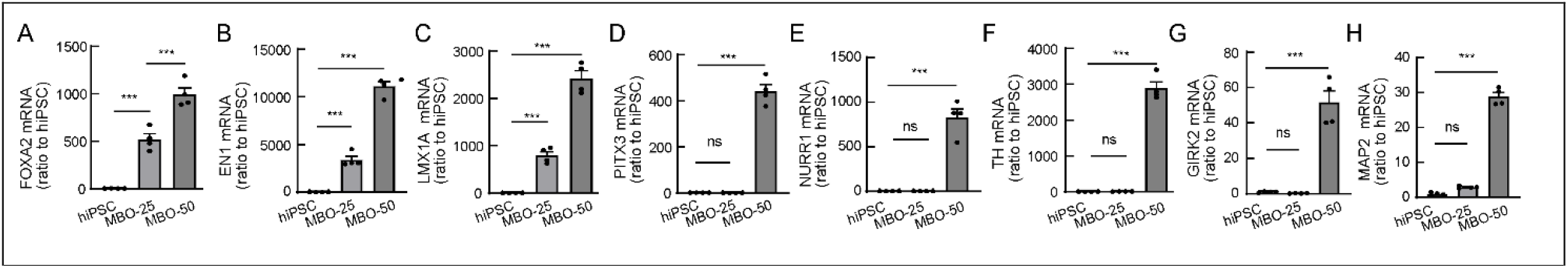
RT-qPCR showed midbrain key markers of brain-region specific patterning on day 25 50. (**A-C**) The elevated expression of midbrain floorplate precursor maker FOXA2, EN1, and LMX1A on day 25 and 50 (n=4). (**D-G**) The elevated expression of more mature marginal zone dopaminergic marker PITX3, NURR1, TH, and GIRK2 on day 50 (n=4). (**H**) The bar graph showed elevated expression of neuronal marker, MAP2 on day 50 (n=4). \*\*\**p*<0.001, one-way ANOVA, n=4.

### 3.4. We specified HPC from hiPSC

KOLF2.1J cell line was subjected to HPC specification using the STEMdiff™ Hematopoietic Kit (STEMCELL Tech.). HPC suspension cells were harvested at 12 days and analyzed by flow cytometry with fluorophore-conjugated antibodies against the HPC surface markers proteins CD43-APC, CD45-FITC, and CD34-PE antibodies. We observed the fraction of CD43^+^ cells to be 94.3% and CD45^+^ CD34^+^ cells to be 24.2%. A high percentage of CD43^+^ cells (> 90%) confirmed the efficient production of HPCs (**Figure 4**).

**Figure 4.**
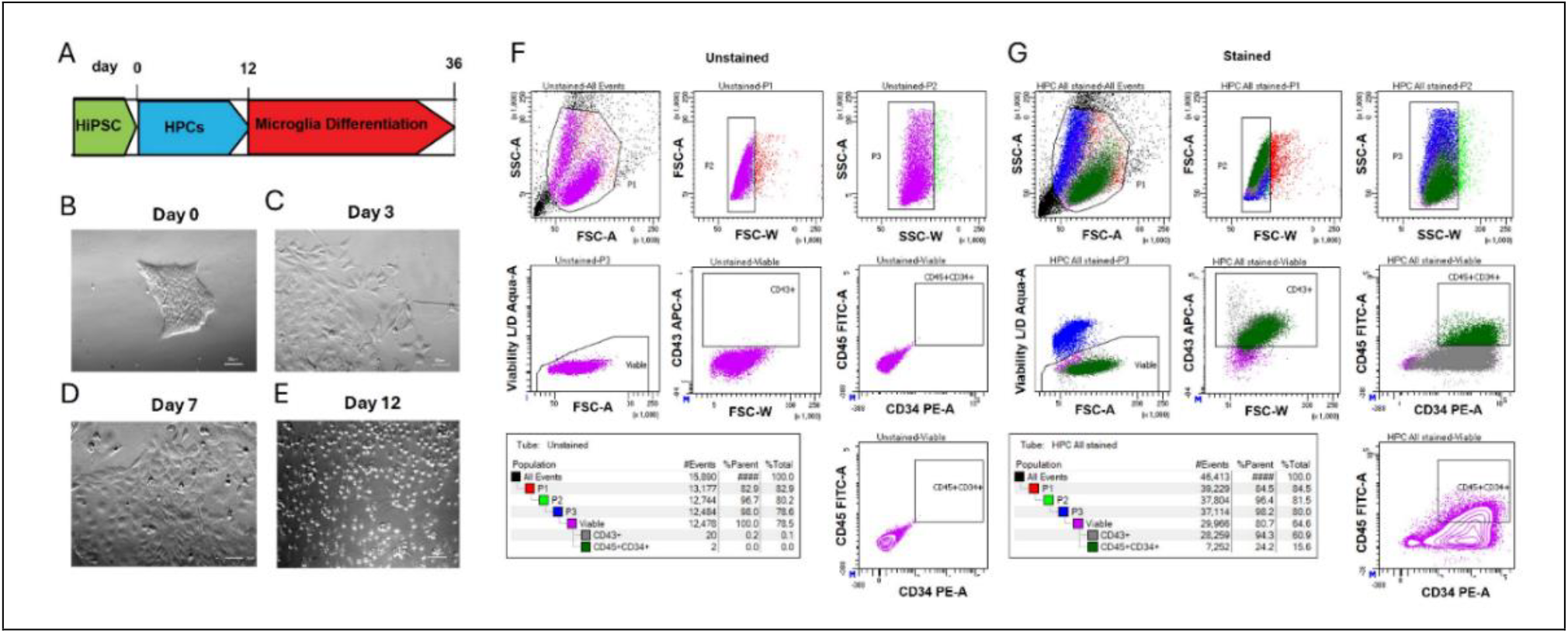
Human iPSCs differentiated into hematopoietic progenitor cells (HPCs). (**A**) The time course of differentiation of microglia from HPCs. (**B-E**) Representative images of HPCs at different time points (bright field), scale bar, 50um. Representative flow cytometry plots for (**F**) unstained HPCs(control) and for (**G**) stained HPCs (CD43^+^ and CD45^+^ CD34^+^) are shown.

### 3.5. Microglia were generated from HPCs

We generated microglia using iPSC-derived HPCs and following the STEMdiff microglia differentiation kit (STEMCELL Technologies Inc.). After 24 days of microglia differentiation and 2 days of maturation, 2 x 10^4^ cells were analyzed by immunohistochemistry using antibodies against the microglia marker proteins Iba1, CD11b(OX-42), and CD45-FITC and counterstained with DAPI (see materials and methods). Fluorescence images were captured using a fluorescent microscope with a 20x objective. Four images were randomly selected for counting CD11b^+^ CD45^+^ cells or Iba1^+^ cells with DAPI, and the percentage of positive cells calculated as a fraction of total DAPI^+^ cells. A high percentage of CD11b^+^CD45^+^ cells (>96%) and Iba1^+^ cells (>90%) (**Figure 5**) indicated that microglia were successfully differentiated from HPCs and could be used towards integration into midbrain organoids.

**Figure 5.**
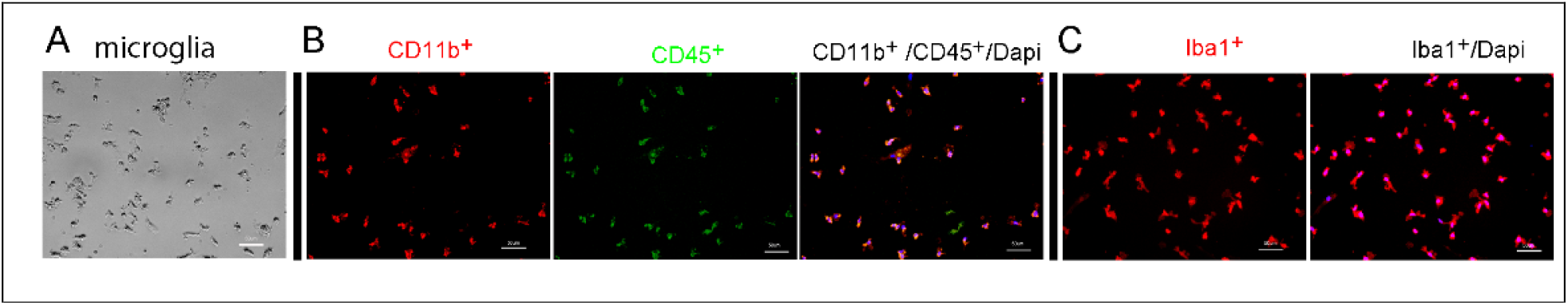
Representative images of microglia differentiated from HPCs. (**A**) Bright field representative image of microglia. (**B-C**) Representative images of microglia immunostaining with microglia marker CD11b^+^CD45^+^(**B**) and Iba1^+^ (**C**). CD11b^+^CD45^+^ cells with Dapi and Iba1^+^ cells with Dapi were counted and averaged out to total number of cells, scale bar = 50µm.

### 3.6. Microglia were integrated into midbrain organoids in a co-culture system

On day 80 of midbrain organoid, differentiated microglia (2.5x10^5^ cells) were applied to each midbrain organoid with co-culture medium, containing organoid maintenance medium (cat#100-0120, STEMCELL Tech.) and microglia supplement 2 (cat#100-0023, STEMCELL Tech.). Co-culturing was continued for 7 days. Then we confirmed the condition of microglia integration into midbrain organoids by immunohistochemistry with Iba1, GFAP, NeuN, MAP2 (another neuronal marker), TH (dopaminergic cell marker), and MnSOD expression at Day 88 (**Figure 6**). These findings confirmed that microglia efficiently populated into midbrain organoids to produce immune-competent microglia-integrated midbrain organoids (MIO).

**Figure 6.**
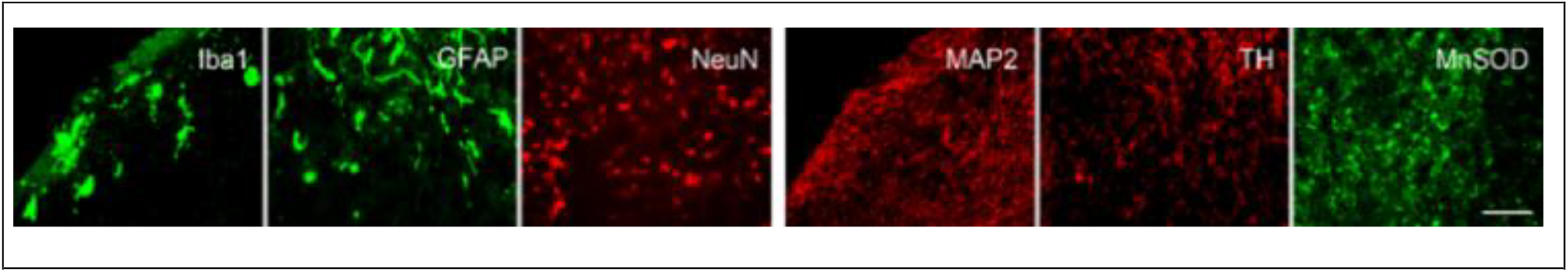
The immunohistochemistry of Iba1 (microglia marker), GFAP (astrocyte marker), NeuN and MAP2 (neuronal markers), TH (dopaminergic marker), and MnSOD expression in midbrain-integrated organoids at day 88, scale bar, 50um.

### 3.7. Microglia-integrated organoids (MIO) model responded to morphine withdrawal

To examine the effect of persistent morphine exposure to MIO, we treated MIO with increasing concentrations of morphine (day1,10 µm; day 2, 20 µm; day 3, 30 µm; day 4, 40 µm, and day 5, 50 µm) each day. After 5 days of morphine treatment, fresh co-culture medium without morphine was replaced for an additional 7 days (morphine withdrawal). The MIO were then collected for neurochemical analysis. We found that morphine withdrawal increased TNFα gene expression and lowered Sirt3 and MnSOD gene expression using RT-qPCR (**Figure 7A-C**).

**Figure 7.**
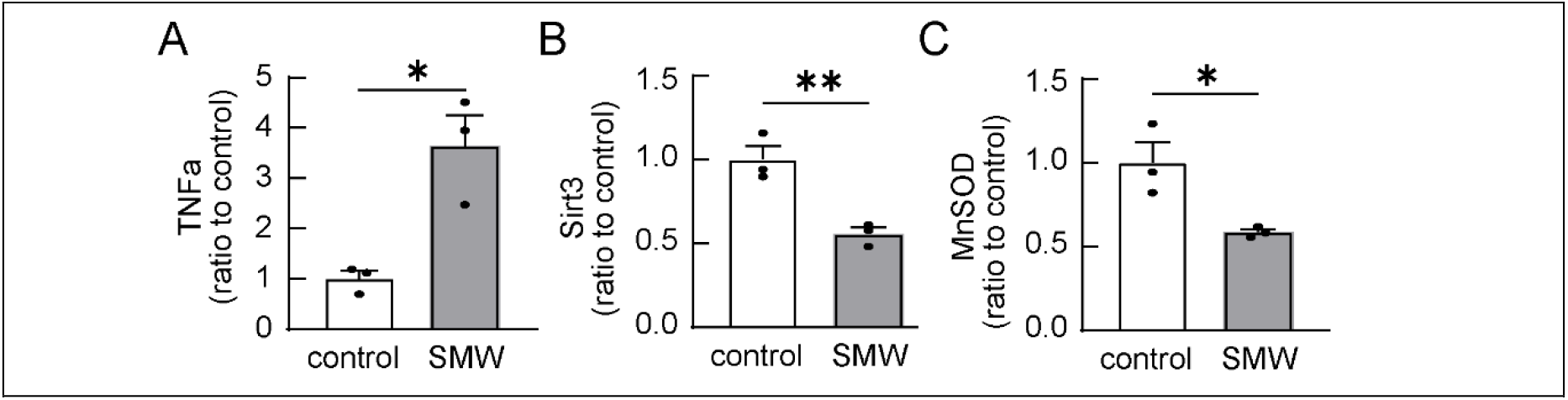
RT-qPCR showed that SMW increased (**A**)TNFα gene expression, but decreased (**B**) Sirt3 and (**C**) MnSOD gene expressions. \**p*<0.05, \*\**p*<0.01, *t* test, n=3.

## 4. Discussion

In this pilot study, using hiPSC KOLF2.1J cell line, we developed our approach for generating both midbrain organoid and microglia, and verified that our midbrain organoid was successfully integrated with microglia. Chronic morphine exposure upregulated TNFα, and downregulated Sirt3 and MnSOD in MIO.

Human iPSC-derived organoid as a self-organized 3D tissue, can mimic the key functional, structural and biological complexity of an organ.^9, 10^ Using neural organoids one can bridge the knowledge gap between animal models and human studies in many neurological disorders.^13-21^ Brain organoids differentiated from human iPSCs to small brain-like structures provide an unprecedented opportunity to elucidate the exact mechanisms of human diseases.^10, 11^ Based on the human origin, the brain organoids likely better match the genomic and structural features compared to animal models.^19, 21-24^ Fernandes and colleagues confirmed the expression of opioid receptors within the iPSC-derived cerebral organoids treated with opioids, and presented higher dopamine secretion recapitulating an important physiological event after opioid exposure.^37^ The Haddad group used iPSC-derived organoid model with prenatal opioid exposure^38-42^ to study the neurobiology of OUD. Notaras and colleagues exposed the forebrain organoids to a panel of narcotic and neuropsychiatric-risk factors and analyzed downstream alterations in transcription, proteomics, and metabolomics.^25^ Ho and colleagues reported that iPSC-derived forebrain organoids treated with opioids displayed a significant influence on transcription regulation in glial cells.^26^

Microglia are the most abundant mononuclear phagocytes in the central nervous system (CNS).^43, 44^ During brain development, microglia establish contacts with neural progenitors to support neurogenesis and proliferation.^45^ Microglia are differentiated from the mesodermal pial cells that invaded the brain during embryonic development,^46, 47^ however other cells are derived from the neuroectodermal lineage.^47^ Microglia play important role in chronic morphine use.^6, 7, 48, 49^ It is reported that microglia can innately develop within a cerebral organoid brain microglia-integrated organoid and display their characteristic ramified morphology ^47^. A report showed an approach to deriving microglia from human iPSCs and integrating them into midbrain organoids to study brain neuroinflammation.^50^ Here, using KOLF2.1J cell line, we developed our approach to generating both microglia and midbrain organoid and verified that our midbrain organoid was successfully integrated with microglia.

The cellular and molecular mechanisms of drug abuse in the brain are extremely complex, targeting multiple neurotransmitter/neuropeptide systems. Chronic morphine use can activate glia and induce neuroinflammation.^6-8^ Glia equipped with several toll-like receptors (TLRs), are implicated in pathological processes of many neurological diseases.^51, 52^ Opioids, such as morphine, oxycodone, and remifentanil, can bind to MD2 and activate TLR4 signaling,^53, 54^ and transcriptional factor nuclear factor-κB (NF-κB) ^55^, leading to the production of proinflammatory molecules. Our preliminary data showed that morphine dependence increased TLR4 and NF-kB protein expression in the midbrain ventrolateral periaqueductal gray in mice (*data not shown*). Here, we found that morphine withdrawal induced upregulation of classical proinflammatory factor TNFα in the MIO.

Mitochondria play a crucial role in producing adenosine triphosphate. The processes also lead to the release of reactive oxygen species (ROS) that may cause oxidative stress.^56^ Chronic opioid use increases product of ROS and decreases the function of enzymatic antioxidants such as superoxide dismutase, catalase, and glutathione peroxidase.^57-59^ ROS is considered the main cause for cellular damage, in turn resulting in higher ROS formation, leading up to a “vicious cycle”.^60, 61^ Oxidative stress resulting from mitochondrial dysfunction contributes to chronic inflammation.^59, 62^ It is reported that inflammatory cytokine TNFα induces Sirt3 impairment, increases SOD2 acetylation and enhances mitochondrial 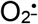 in human endothelial cells.^63^ Our previous study showed that TNFα induced mitochondrial oxidative stress in neuropathic pain states.^36, 64^ Opiates such as heroin and morphine are able to induce the formation ROS particularly superoxide in cultured cells.^65, 66^ Antioxidants inhibit the oxidative stress and opioid dependence in animals.^67, 68^ SOD scavenger enzymes convert superoxide radicals into H_2_O_2_ and molecular oxygen. The steady-state level of superoxide is inversely proportional to the activity of MnSOD located in the mitochondrial matrix that appears to be a central player in the redox biology of cells and tissues.^69^ Heroin decreased the total antioxidant capacity of serum and the antioxidative enzymes activities in brains.^70^ Mitochondrial Sirt3 regulates every major aspect of mitochondrial biology, including ROS detoxification and mitochondrial dynamics.^71-73^ Sirt3 can rescue neuronal loss in various neurodegenerative models.^74^ Our preliminary studies demonstrate that chronic morphine use decreased expression of Sirt3 in the midbrain (*data not shown*). Here, we found that chronic morphine treatment to MIO lowered expression of MnSOD and Sirt3, which in line with the previous reports above.

## 5. Conclusion

Newly emerged human iPSC-derived midbrain MIO can at least partially mimic the functional, structural and biological complexity of brain. Midbrain MIO provides important approach to examining the mechanisms of neuroinflammation and neurodegeneration. It can also be utilized as an effective tool for drug screening and toxicity testing.

## Acknowledgement

We thank the support from the Department of Anesthesiology, University of Miami, FL. This work was partially supported by grants from the NIH 5R01DA034749 (SH), NIH R01DA047157 (SH) and VA 5I01BX005114 (SH). This research partially received funding from the Office of the Executive Dean for Research at the University of Miami Miller School of Medicine (NEURO-TSA-2024-2, ZZ) and the US Department of Defense (AL240243, ZZ). Authors acknowledge the Flow Cytometry Shared Resource (FCSR) of the Sylvester Comprehensive Cancer Center at the University of Miami (RRID: SCR022501) supported by NIH P30CA240139 for technical assistance. Authors also acknowledge Dr. Xue Zhong Liu (Otolaryngology) and Dr. Derek Dykxhoorn (iPSC Core, HIHG) at the University of Miami for technical assistance.

